# Peripheral blood exosomes pass blood-brain-barrier and induce glial cell activation

**DOI:** 10.1101/471409

**Authors:** Diana M. Morales-Prieto, Milan Stojiljkovic, Celia Diezel, Priska-Elisabeth Streicher, Franziska Röstel, Julia Lindner, Sebastian Weis, Christian Schmeer, Manja Marz

**Author notes:** these authors contributed equally to this work (Diana M. Morales-Prieto, Milan Stojiljkovic, Christian Schmeer, Manja Marz).

## Abstract

**Background:** Exosomes are involved in intracellular communication and contain proteins, mRNAs, miRNAs, and signaling molecules. Exosomes were shown to act as neuroinflammatory mediators involved in neurodegenerative diseases including Alzheimer’s disease (AD), Parkinson’s disease (PD), amyotrophic lateral sclerosis (ALS). Brain aging has been associated to increased neuroinflammation. In addition, a decreased extracellular vesicle concentration was observed in aging tissues. The specific mechanisms how exosomes and aging are connected are not known yet.

**Results:** Here we have shown that peripheral injection had almost no effect on selected gene expression in the liver. However, exosome injection has led to changes in the specific markers of glial cell activation (*CD68, Iba1, GFAP*). Interestingly, only injection of exosomes isolated from aged mice induced significant activation of astrocyte cells, as shown by increased *GFAP* expression.

**Conclusion:** Transcription levels of genes *GFAP, TGF-β, CD68, Iba1* known to be involved in glial cell function are significantly changing after introduction of peripheral extracellular vesicles. Exosomes were able to pass blood brain barrier and induce glial cell activation. *GFAP* known to be a specific astrocyte activation marker was significantly higher expressed after injection of old but not young exosomes, indicating a possible role of exosomes in the mechanisms of brain aging.

## Introduction

Exosomes are specialized membranous vesicles (40-100 nm in diameter) of endocytotic origin^19^. Exosomes are formed intracellularly via endocytic invagination and are generated by outward budding at the endosomal membrane of the multivesicular bodies (MVBs)^14^. Exosomes are involved in intracellular communication^8^ and contain proteins, mRNAs, miRNAs, and signaling molecules that reflect the physiological state of their cells^9^.

Recently, experimental evidence was provided suggesting that circulating exosomes may act as a neuroinflammatory mediator in systemic inflammation^5^. Neuroinflammation is a common pathological feature of neurodegenerative diseases including Alzheimer’s disease (AD), Parkinson’s disease (PD), frontotemporal lobar dementia (FTD), and amyotrophic lateral sclerosis (ALS) and is represented by glial activation and pro-inflammatory cytokine production by the central nervous system (CNS)-resident cells^7^. Furthermore, several evidences indicate a critical role of neuroinflammation in brain aging^11^,^12^.

Accumulating evidences implicate extracellular vesicles (EVs) in the aging process: It has been shown that the plasma EV concentration decreases with human age^4^. The same study shows that B cells but not T cells internalize EVs and that B cells from the elderly uptake more EVs. Plasma EVs isolated from young but not from elderly donors promote the osteogenic differentiation of mesenchymal stem cells in a galectin-3 dependent manner^18^. Furthermore, EVs purified from the elderly suppress cell proliferation and osteogenic differentiation of bone marrow stromal cells^3^.

More recently, exosomes secreted by stem/progenitor old (24 months) or young (3 months) animals were cells of the healthy hypothalamus were associated with a slowing of aging^20^. In particular, speed of aging was suggested to be controlled by exosomal miRNAs from the hypothalamic stem cells, further supporting a role of EVs in the aging process. The impact of peripheral blood-derived exosomes on aging, in particular from the CNS has not been assessed yet. Also, putative targets of peripheral exosomes, in particular in the aging brain, still remain to be elucidated.

Here, we evaluated putative target genes of peripheral blood exosomes from aged wild type mice in liver and brain. Interestingly, stained exosomes were able to pass the blood-brain-barrier and had an impact on neuroinflammation and glia cell activity.

Brain aging is characterized by an increase in inflammatory mediators like TNF-*α*, IL-6, TGF-*β* and gliosis induced by glial cell activation. As injection of old blood led to cognitive decline in young mice, we wanted to check whether peripheral injection of purified exosomes could pass the blood brain barrier, be taken up by glial cells and induce a significant transcriptional changes in the brain. Here, we found that a single exosome injection leads to a change of glial cells and induce gene transcription even as soon as 4 h, and continue to increase with time. We found glial cells gene transcriptional activation without changes in the inflammation cascade. Both young and old exosomes induced differential transcription of glial genes. We also found first hints of age specific effect of exosomes. Further studies should investigate and analyze the content of exosomes to discover the mechanism behind this effect.

## Material and Methods

### Animals experiments

C57BL/6J mice, originally obtained from the Jackson Laboratories, were bred and maintained at the animal facility (ZET) of the University Hospital Jena under specific pathogen-free conditions with a 12 h day/nigh cycle and fed ad libidum. All animal experiments were approved by the local regulatory board (Thuringian State Office for Consumer Protection and Food Safety (02-044/16). For exosome isolation, blood from donor collected in citrate collection tubes. Recipient animals were injected with 100 *µ*l of isolated exosomes or vehicle into the tail vein of 11-12 week old mice. Mice were sacrificed 0.5 h, 4 h and 24 h after injection (see Supplementary Tab. S1). Blood was collected in heparinized syringes via cardiac puncture and mice were subsequently perfused with PBS. Brain, liver and other organs were harvested for RNA and localization analysis.

### Exosome isolation, quantification and labeling

Exosomes were isolated from pooled plasma (approx. 480 *µ*l/mouse) by differential ultracentrifugation. Samples were centrifuged at 10,000 g for 10 min and then at 18,900 g for 30 min at 4°C to remove cell debris and microvesicles. Supernatants were filtrated through a 0.22 *µ*m membrane filter and then centrifuged at 4°C at 100,000 g for 5 h using the Beckman Coulter Type 55Ti rotor. Pellets were resuspended in PBS and suspension was centrifuged again at 100,000 g overnight. Exosomes were resuspended in 100 *µ*l of PBS and protein content was quantified by Micro BCA^™^Protein Assay (Thermo Scientific^™^). For the PKH67-staining (PKH67GL-1KT, SIGMA-ALDRICH), exosomes were dissolved in 500 *µ*l of Diluent C and mixed with 500 *µ*l of PHK67 dye diluted in diluent C (1:250 v/v). Staining was stopped after 2 min by addition of 1 % BSA in Isotonic Saline Solution 0.9%. PKH-67 labeled exosomes were centrifuged over night at 100,000 g. Pellet was resuspended in 100 *µ*l saline solution and protein was quantified by Micro BCA^™^Protein Assay. Naive recipient mice received 2 *µ*g of protein equivalents of exosomes.

### Histological Analysis

Recipient mice receiving 2 *µ*g of protein equivalents of exosomes from old donors were euthanized 30 min after injection. After perfusion with PBS, brain and liver tissues were harvested and cryopreserved in sucrose. Tissues were embedded in optimal cutting temperature compound and sectioned on a Leica CM1850 cryostat at a thickness of 20 *µ*m. Sections were attached to microscope slides and allowed to dry on a slide warmer for an hour at 37°C. Sections were observed under an LSM510 microscope (Zeiss).

### EV uptake in primary isolated cells

Primary glial cultures containing ∼80 % astrocytes and 20 % microglia^13^ were grown in a 4-chamber culture slide (FALCON) at 100,000 cells per well and, after washing with PBS, incubated with 20 *µ*l/ml PKH67-labelled EVs (∼400*µ*g/well) for 1 h in DMEM containing ED-FBS. For immunodetection, cells were fixed with 4 % PFA (Paraformaldehyde) for 30 min and permeabilized and blocked with 0.3 % Triton X-100/3 % NDS (normal donkey serum) in PBS for 30 min, washing the cells with PBS between the steps. Primary antibodies, rabbit-anti-Iba1 (WAKO Chemicals, U.S.A.) and goat-anti-GFAP (Abcam), where applied for incubation at 4°C overnight in a dilution of 1:500 in PBS/10 % BSA. After washing with PBS, secondary rhodamin antibodies (Jackson Immunoresearch, U.S.A.) and DAPI (4-6-Diamidino-2-phenylindole, SigmaAldrich) were added in a dilution of 1:500 in PBS/10 % BSA and incubated for 1 h at room temperature. Cells were washed with PBS and ddH2O, covered with mounting medium for fluorescence (VECTASHIELD R) and a cover slip and observed under an AxioPlan2 microscope (AxioObserver Z.1 – Zeiss).

### Analysis of gene expression by qPCR

Mixed glial cultures were seeded at 100 000 cells per well and incubated with EV (∼400 *µ*g/well) for 24 h. Thereafter, RNA was extracted using QIAzol reagent (Qiagen) combined with isopropanol precipitation. The RNA concentration, quality and integrity were determined using a Nanodrop (Thermoscientific, Waltham, MA, USA) and QIAxcell Systems (Qiagen). cDNA was synthesized from 500 ng of RNA/reaction using a RevertAid First Strand cDNA Synthesis kit (Thermoscientific). qPCR was performed using a LightCycler 480 SYBR Green kit (Roche, Germany). Detection and quantification were conducted with a Rotor gene cycler and Rotor gene Q software (Qiagen). The housekeeping genes *Gapdh* and *Hmbs* were used for normalization. The relative gene expression was calculated using the 2-^ΔΔCT^ method^6^. The primers used are listed in Supplementary Tab. S2.

### Statistical analysis

Data are presented as the mean ±SEM, and *n* represents the number of independent experiments. One-way analysis of variance (ANOVA) with Holm-Sidak correction or Student’s t test were used for analysis. Statistical analysis was performed using Sigma plot version 12.5 (Systat, San Jose, CA, USA). P-values of 0.05 or less were considered significant.

## Results

### Localization of peripheral injected exosomes in recipient mice

Isolated exosomes from aged mice (24 months old) were stained with PKH67 and injected into the tail vein of 3-month-old mice for 30 min. Sections of brain and liver tissue were counter-stained with DAPI for localization of cell nuclei and observed under a fluorescence microscope. PKH67-labeled exosomes (green fluorescence) were detected in brain but not in liver tissue (Fig. 2). Distribution of exosomes in brain tissue was not homogenic as observed by the agglomeration of green signals in some areas of the tissue. A specific pattern demonstrating the presence of exosomes on specific areas of the brain could not be determined.

**Figure 1:**
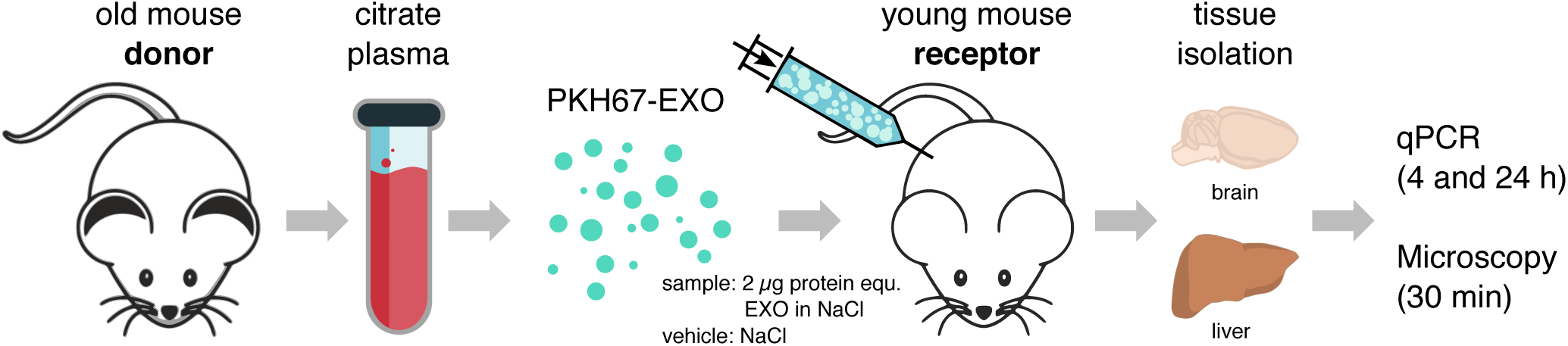
Experimental setup. Exosomes are extracted from blood of old mice, PKH67-stained and in young mice injected. Brain and liver were isolated for histopathological analysis and for qPCR of specific genes

**Figure 2:**
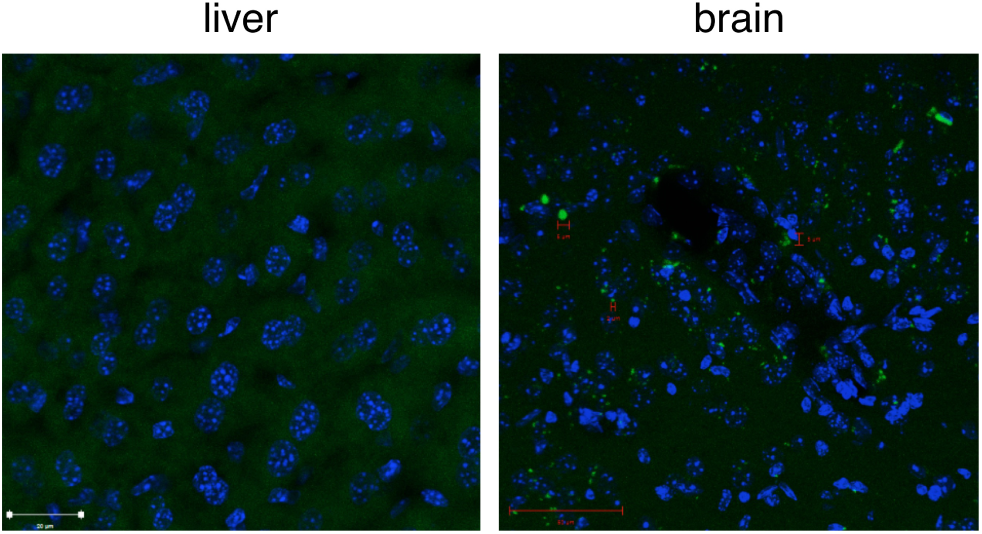
Representative micrographies of brain and liver 30 min after injection of PKH67-stained exosomes into tail vein. Blue – DAPI, green – PKH67-stained exosomes.

### Effect of aged exosomes on gene expression in liver tissue

Young mice were injected with PKH67-labeled exosomes from aged mice (24 months old) or vehicle (0.1 % BSA in PBS) and sacrificed after 4 h or 24 h. We extracted total RNA from liver and performed qPCR analysis for some the most com-mon activation/inflammatory genes: *TNF-α, Iba1, CD68, TGF-β, IL-10, IL-6, iNOS*. Compared to vehicle, peripheral exosome injection significantly decreased *IL-6* at 24 h. *IL-10* was significantly increased in treated mice after 24 h compared to 4 h but this effect was not different to the observed with the vehicle control (see Fig. 3).

**Figure 3:**
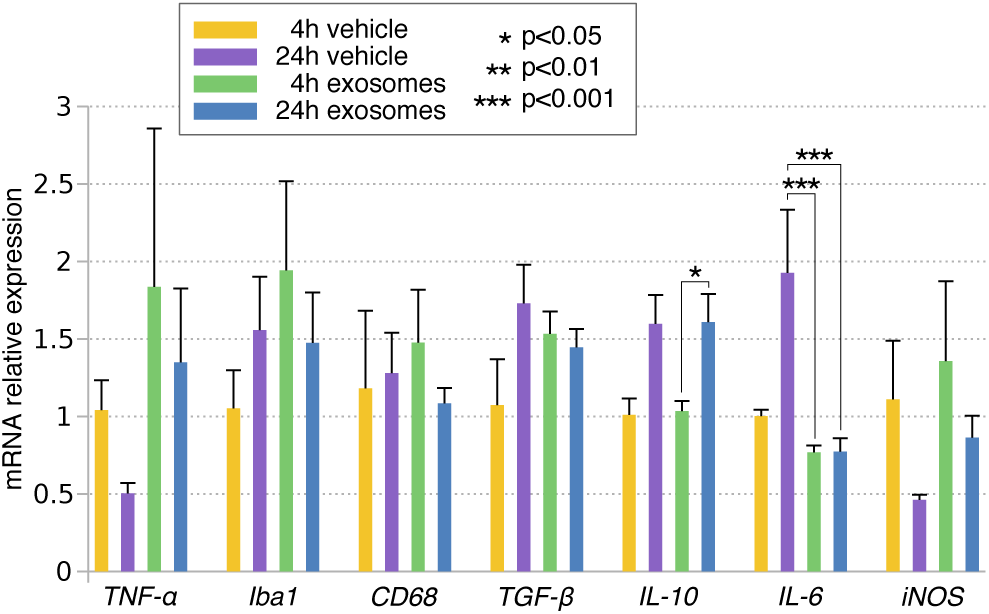
mRNA analysis of liver tissue after trea™ent with aged exosomes (4 h, 24 h). *p<0.05, ***p<0.001, One way ANOVA with Holm-Sidak correction.

### Aged exosomes alter gene expression in brain tissue *in vivo*

Using the same experimental design described in the previous section, we analyzed activation/inflammatory genes in brain tissue of young mice injected with aged exosomes. We observed alteration of genes related with activation (*GFAP, Iba1, CD68, TGF-β*) but no with brain inflammation (*TNF-α, IL-6,IL-1β*) when compared to non-injected mice (see Fig. 4 and Supplementary Fig. S1). To discard the possible effects caused by the injection procedure or the vehicle itself (0.1 % BSA), animals injected with vehicle were included as controls.

**Figure 4:**
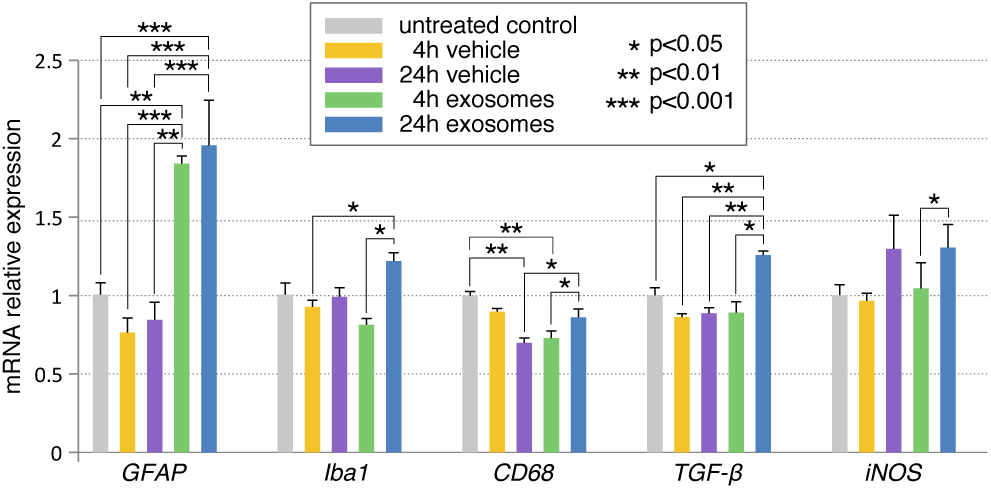
Peripheral exosomes induce gene expression in brain tissue. Mice were injected with aged exosomes or vehicle (0.1 % BSA) and sacrificed after 4 h or 24 h. *p<0.05, **p<0.01, ***p<0.001, One way ANOVA with Holm-Sidak correction.

We could clearly show that the vehicle did not have significant impact on gene expression and that the effect of exosome induction of glial cell activation was reproducible, as observed by the significant increase of *GFAP* (4 and 24 h), *Iba1* (24 h) and *TGF-β* (24 h) transcription (see Fig. 4).

### Differential effect of aged and young exosomes on brain gene expression

To evaluate whether aging of donor mice could modulate the observed effects in recipient mice, expression pattern in the brain was compared 24 h after injecting exosomes from young and old mice. A similar induction pattern for *Iba1, CD68, TGF-β* and *iNOS* was observed after injection of young and aged exosomes compared to untreated controls. Remarkably, elevated transcription ∼14 % was observed in *GFAP* after injection of aged compared to young exosomes resulting in significance of only aged exosomes compared to untreated controls (Fig. 5).

**Figure 5:**
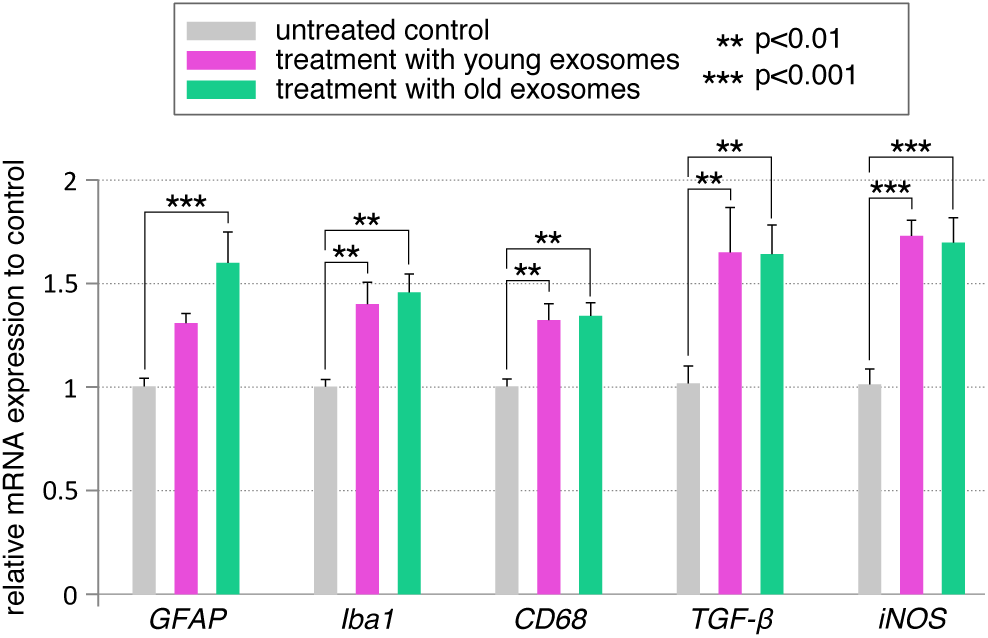
Differential effect of aged and young exosomes on brain gene expression. Circulating exo-somes were isolated from aged and young donor mice. Recipient mice were injected with 2 *µ*g of exosomes or vehicle and sacrificed after 24 h. **p<0.01, ***p<0.001, One way ANOVA with Holm-Sidak correction.

### Uptake of peripheral exosomes by microglia and astrocyte cells *in vitro*

To determine whether a specific cell population of glia cells could be targeted by peripheral exosomes, an uptake study was designed using a primary mixed glial culture containing 80 % astrocytes and 20 % microglia treated with PKH-labeled aged exosomes. Astrocytes and microglia were immunostained with specific markers (*GFAP* and *Iba1*, respectively), nuclei were counter-stained with DAPI and colocalization with exosomes (green) was investigated by fluorescence microscopy. Preliminary results demonstrate that exosomes interact with both populations as observed by the presence of green dots and aggregations throughout the slides. Exosomes were localized mostly in the cytoplasm compar™ent and overlap in major extend with the signals of *Iba1* indicating microglia as the major recipient cells (see Fig. 6).

**Figure 6:**
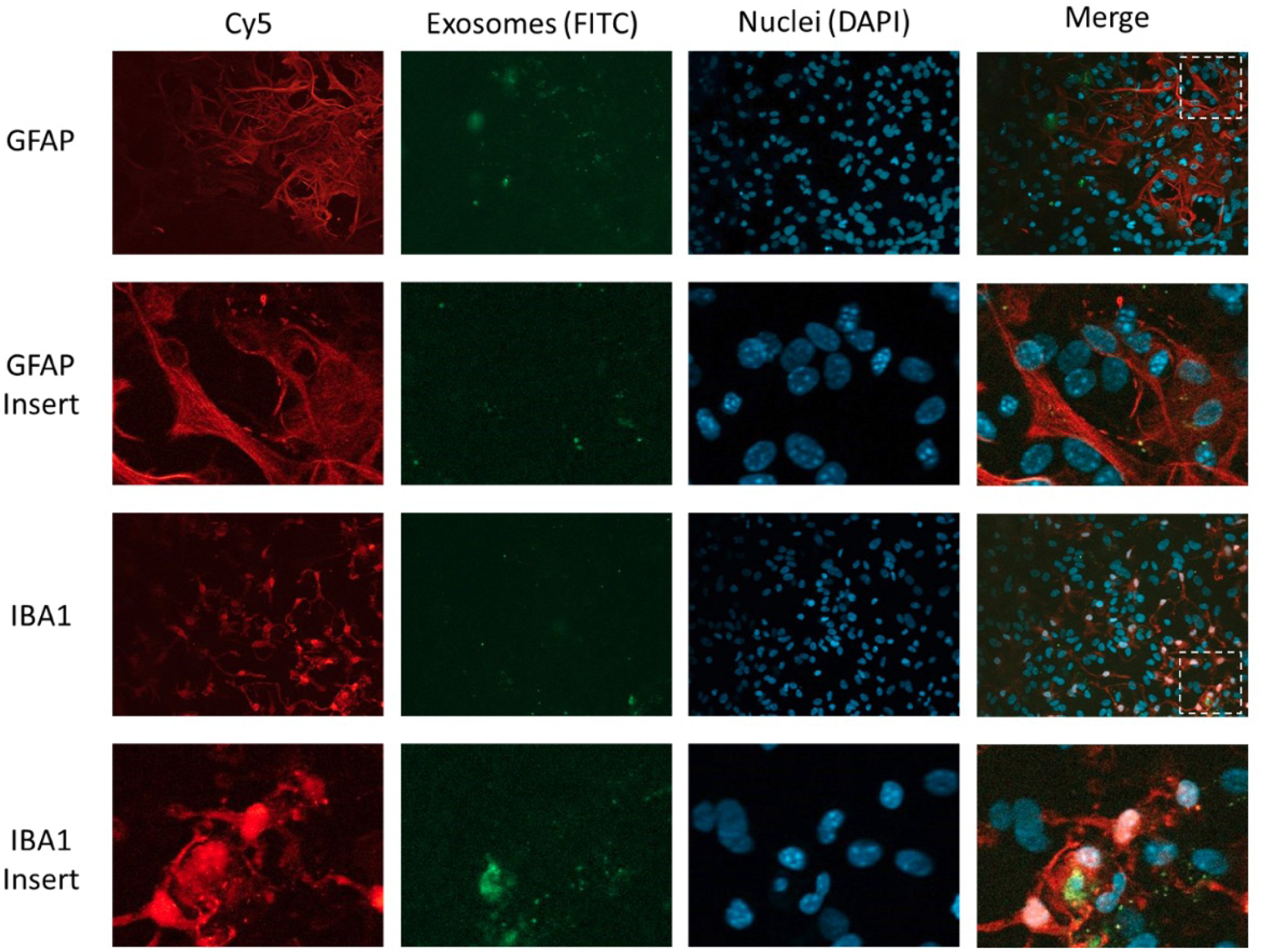
Uptake of aged exosomes by astrocytes and microglia cells *in vitro*. Primary mixed glial culture containing astrocytes and microglia was incubated with PKH67-labeled exosomes for 1 h. Cells were stained with anti-Cy5 antibodies against GFAP (astrocytes) or Iba1 (microglia), nuclei was counter-stained with DAPI. Exosomes are taken up by both cell population but co-localization with microglia cells was more frequent.

### Effect of aged and young exosomes on gene expression *in vitro*

To further investigate the differential effects by aged versus young exosomes, we used the *in vitro* model of primary mixed cultures described before and investigated gene expression of *Iba1, GFAP, CD68, TGF-β* and *iNOS* after 24 h trea™ent with exogenous exosomes. Using this setting, only *TGF-β* was significantly increased after trea™ent with exosomes from old compared to young donors (Fig. 7).

**Figure 7:**
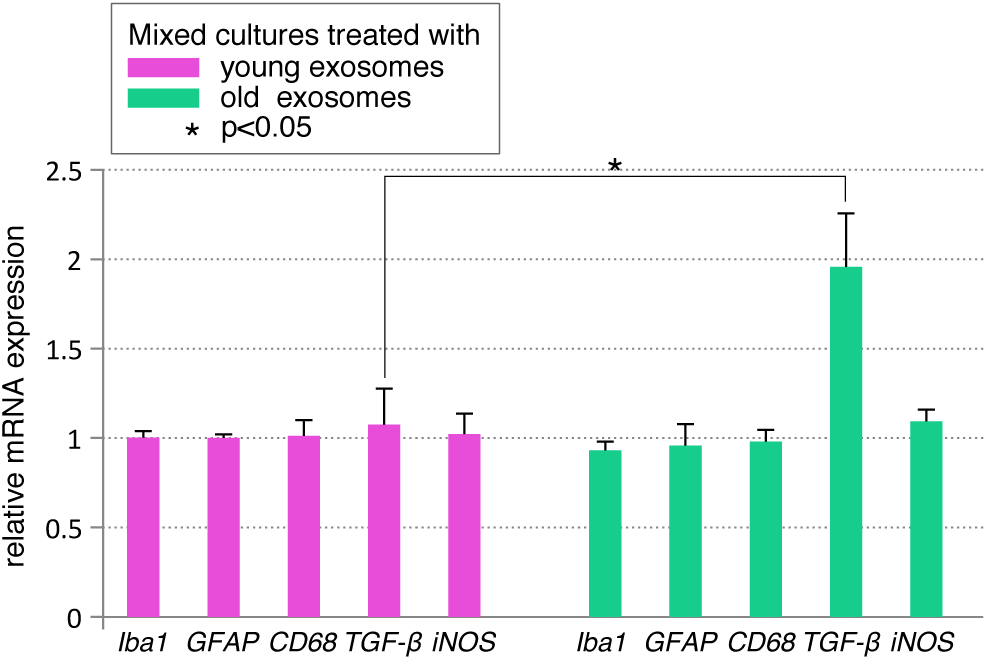
mRNA analysis of mixed cell culture (80 %astrocytes, 20 % microglia) after 24 h trea™ent with exosomes from young and old donor

## Discussion

Exosomes are nowadays recognized as major players in the intercellular communication due to their ability to transfer proteins and genetic information horizontally. Accumulating evidence demonstrates involvement of exosomes in spreading inflammatory signals and oxidative stress in a plethora of human pathologies. For instance, misfolded proteins, disease-associated particles such as *α*-synuclein, A*β*, and prions can be transferred from origin to recipient cells via exosomes driving progression of neurodegenerative disorders including Alzheimer’s disease^2^.

Due to their small size, exosomes can cross the blood-brain barrier and therefore be used as delivery vehicles of drugs into the CNS. At the same time, this ability could result in negative effects when elevated levels of exosomes from the periphery access the parenchyma or when the content of these exosomes deviate from the normal.

Here, we demonstrate that injected exosomes localized in the brain rather than in liver tissue of recipient mice. Whether this effect is due to a specific recognition, accumulation of peripheral exosomes in the small capillary of the brain or a slower clearance in the brain compared to the liver could not be determined with our settings. However, it demonstrates the pivotal role of exosomes in accessing the CNS. Likewise, and in concordance with the localization results, a clear change of transcription signature was observed in brain but not in liver.

Only slight reduction in liver IL-6 and increase in IL-10 levels was found in mice injected with exosomes, which suggests a trend to a reduced inflammatory response. On the other hand, we found major changes in expression of genes related with glia activation (*GFAP, Iba1, CD68, TGF-β*) but not with inflammation (*TNF-α, IL-6, IL-*in brain tissue. These results are in agreement with a previous report showing microglia and astrocyte activation in mice after systemic delivery of serum exosomes from LPS-challenged donors^5^. Recently, several groups have shown that glial cell uptake and release of exosomes may play a role in neurodegenerative diseases^1^. Additionally, glial cells could take up the substances released in blood from the gut microbiome, so called gut-brain axis, which could play a role in glial cell development and activation^15^. Here, we have shown *in vivo* that glial cells can be activated by interaction and internalization of peripheral injected exosomes.

Most strikingly, a significant increase in *GFAP* transcription was observed after injection of exosomes from old but not young donor mice. It was shown previously that aging and rejuvenation are transferable via blood between two mice: In a landmark study from 2011, it was shown that connecting blood vessel systems of a young and an old mouse caused the young animal to age significantly faster^16^. *Vice versa*, in 2014, the same group showed that young blood even reversed the impairments in cognitive function and synaptic plasticity of the older mouse^17^.

To further evaluate whether peripheral exosomes could be specifically recognized by different glia populations, we also used an *in vitro* model consisting of mixed astrocytes and microglia cells. Uptake analysis demonstrates that both populations interact with and take up exosomes within a short period of time, which then co-localize more often with microglia. This may be explained by the phagocytosis ability of microglia cells, which agrees with some studies demonstrating high recognition and internalization of exosomes by macrophages populations in brain^21^ and spleen^10^. In this setting, almost no significant changes in gene expression were found, contrasting with the results in the *in vivo* model. The discrepancy with the findings could be due to a necessary interaction between peripheral exosomes and the blood-brain-barrier causing an indirect activation of glia cells, but also to an already activated state of microglia and astrocytes in the *in vitro* assay induced by the isolation procedure. Compared to young donors, exosomes from old donors resulted in increased *TGF-β* expression in the primary mixed culture suggesting differences in the surface or content of aged exosomes.

Altogether, the experiments shown here indicate that peripheral-injected exosomes selectively target brain tissue and induce activation of glia cells in a mouse model. Furthermore, expression of *GFAP* and *TGF-β* were significantly altered only after trea™ent with aged exosomes suggesting differences in the surface and content of exosomes mediated by aging. These alterations in exosome content could be involved in transferring of the aging phenotype via blood from old to young mice.

## Supplementary Information

**Supplementary Table S1:** Experimental setup for the *in vivo* mouse experiments. Tribe: C57BL/6J; Exo – Murine Exosomes; Veh – NaCl Vehicle; Dea – death post infection; Don – Donor being o. – old or y. – young; His – used for histological analysis.

| Acceptor mouse |  |  |  |  | Experiment |  |  |  |  | qPCR | His |
| --- | --- | --- | --- | --- | --- | --- | --- | --- | --- | --- | --- |
| ID | sex | age<br>m | weight<br>g | t<br>°C | Don | Stain | Exo<br>V<br>μ | Vec<br>V<br>μl | Dea<br>h |  |  |
| 1 | f | 3 | 26.78 | 32.9 | o. | yes | 100 |  | 24 | no | yes |
| 2 | f | 3 | 19.77 | 33.1 | o. | yes | 100 |  | 24 | no | yes |
| 3 | f | 3 | 24.12 | 32.7 | o. | yes | 100 |  | 24 | no | yes |
| 4 | m | 3 | 23.07 | 33.4 | o. | yes | 100 |  | 4 | no | yes |
| 5 | m | 3 | 25.53 | 33.2 | o. | yes | 100 |  | 4 | no | yes |
| 6 | m | 3 | 27.61 | 33.5 | o. | yes | 50 |  | 4 | no | yes |
| 7 | m | 3 | 28.41 | 32.5 | o. | no |  | 100 | 4 | no | yes |
| 8 | f | 3 | 21.05 | 32.9 | o. | no |  | 100 | 4 | no | yes |
| 9 | m | 3 | 27.03 | 33.5 | o. | no |  | 100 | 4 | no | yes |
| 10 | m | 3 | 29.43 | 31.8 | o. | no |  | 100 | 24 | no | yes |
| 11 | f | 3 | 23.29 | 33.2 | o. | no |  | 100 | 24 | no | yes |
| 12 | f | 3 | 22.77 | 33.1 | o. | no |  | 100 | 24 | no | yes |
| 13 | m | 3 | 26.33 | 31.9 | o. | no | 100 |  | 0.5 | no | yes |
| 14 | f | 3 | 22.06 | 32.9 | o. | yes | 100 |  | 0.5 | no | yes |
| 15 | m | 3 | 27.90 | 33.3 | o. | yes | 100 |  | 0.5 | no | yes |
| 16 | m | 3 | 28.23 | 33.0 | o. | yes | 50 |  | 4 | no | yes |
| 17 | f | 3 | 22.67 | 33.3 | o. | yes | 100 |  | 4 | no | yes |
| 18 | m | 3 | 28.84 | 32.4 | o. | yes | 100 |  | 4 | no | yes |
| 19 | m | 3 | 27.03 | 32.9 | o. | yes | 100 |  | 24 | no | yes |
| 20 | f | 3 | 21.58 | 33.2 | o. | yes | 100 |  | 24 | no | yes |
| 21 | f | 3 | 20.23 | 33.3 | o. | yes | 100 |  | 24 | no | yes |
| 22 | m | 3 | 24.67 | 32.8 | o. | yes | 100 |  | 0.5 | no | yes |
| 23 | m | 3 | 25.07 | 32.9 | o. | yes | 100 |  | 0.5 | no | yes |
| 24 | m | 3 | 24.96 | 32.3 | o. | yes | 100 |  | 24 | no | yes |
| 25 | m | 3 | 27.72 | 32.9 | o. | no | 100 |  | 24 | yes | no |
| 26 | m | 3 | 28.99 | 32.6 | o. | no | 100 |  | 24 | yes | no |
| 27 | m | 3 | 25.98 | 32.8 | o. | no | 100 |  | 24 | yes | no |
| 28 | m | 3 | 26.22 | 32.4 | o. | no | 100 |  | 24 | yes | no |
| 29 | m | 3 | 29.12 | 31.8 | y. | no | 100 |  | 24 | yes | no |
| 30 | m | 3 | 26.87 | 31.9 | y. | no | 100 |  | 24 | yes | no |
| 31 | m | 3 | 25.93 | 32.1 | y. | no | 100 |  | 24 | yes | no |
| 32 | m | 3 | 27.15 | 32.5 | y. | no | 100 |  | 24 | yes | no |

**Supplementary Table S2:**
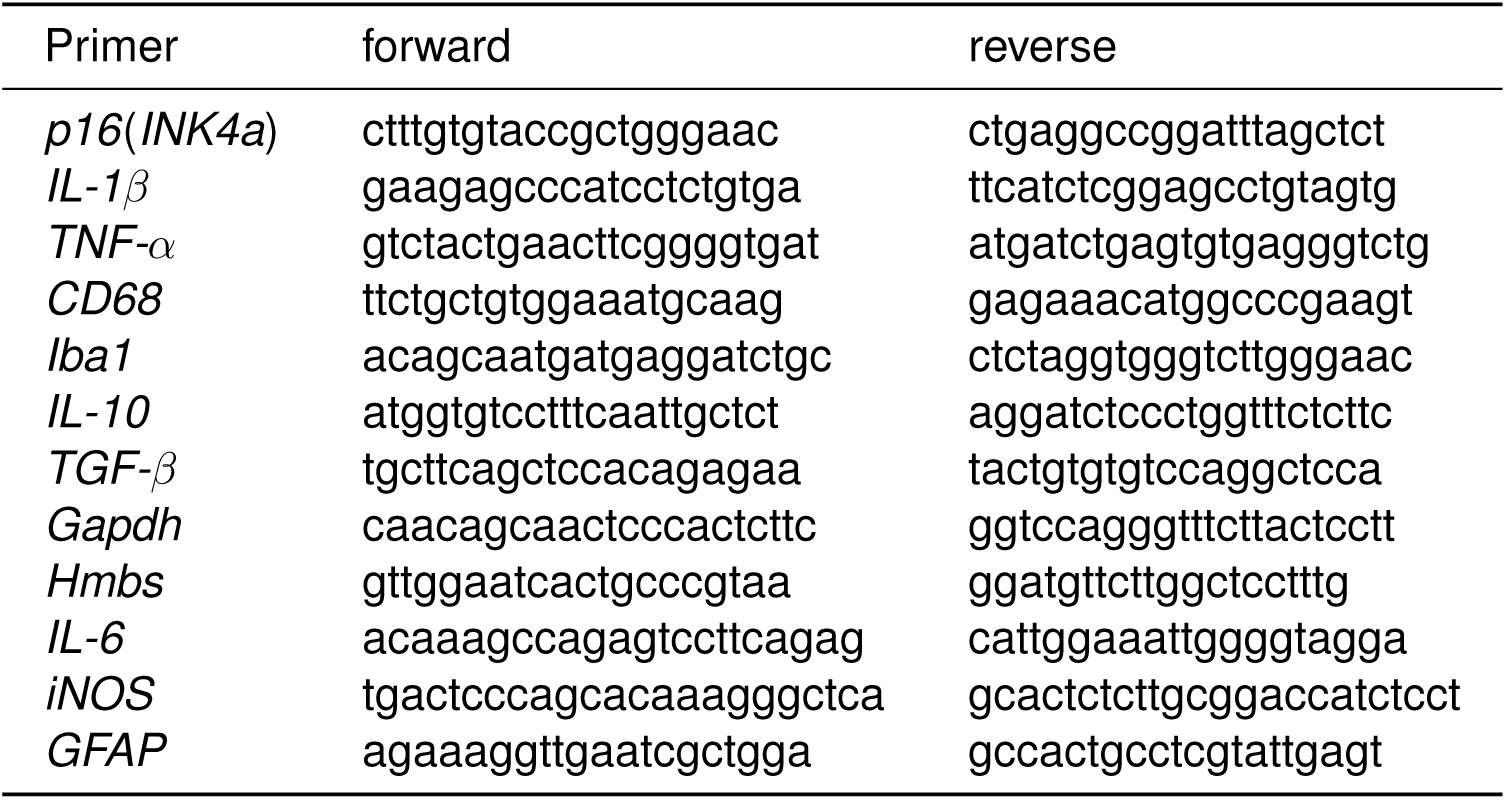
Primer sequences used for quantitative RT-PCR.

**Supplementary Figure S1:**
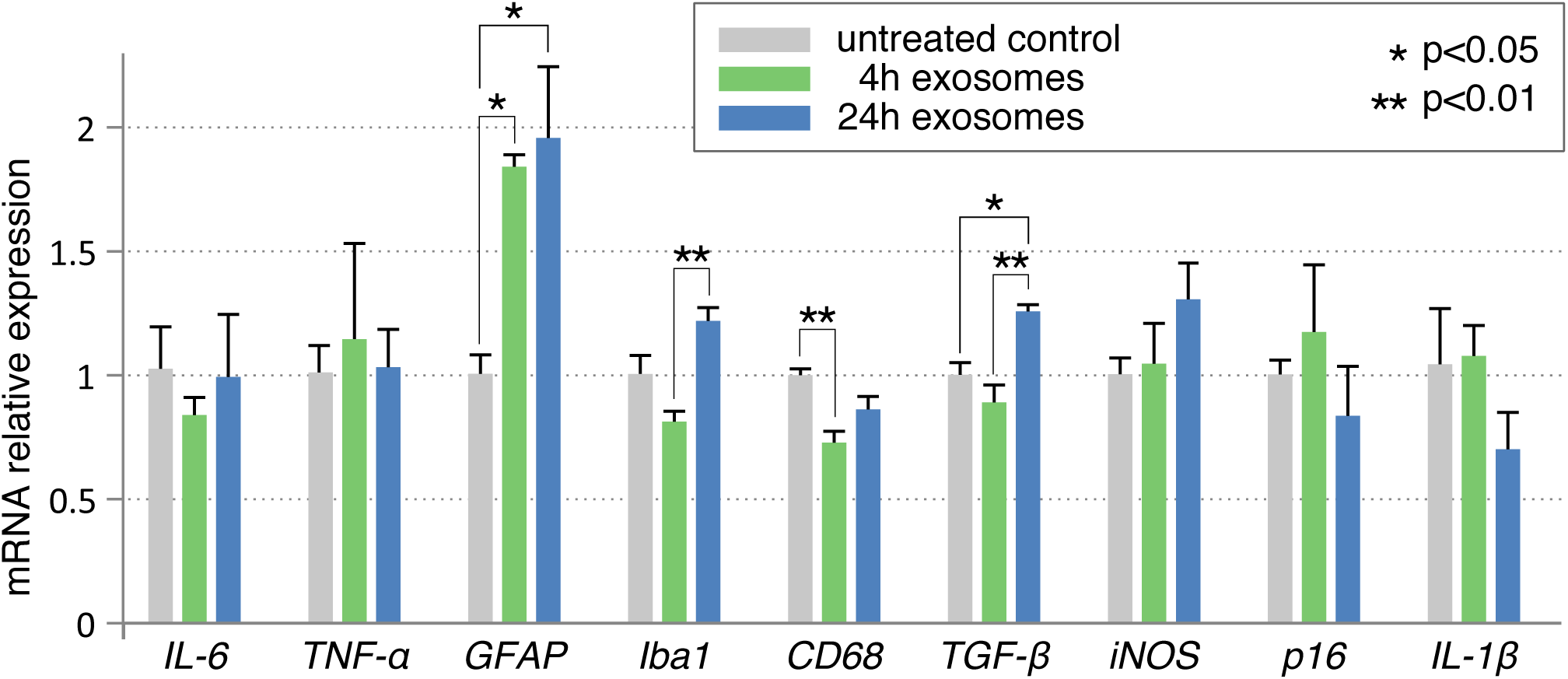
mRNA Analysis of the whole brain tissue after exosome trea™ent (4 h, 24 h). Four out of nine genes from brain samples are tested via qPCR for a difference after injection of exosomes of old mice. *GFAP, Iba1, CD68* and *TGF-β* show significant changes after exosome trea™ent. *p<0.05, **p<0.01, One way ANOVA with Holm-Sidak correction.

